# The vertical position of visual information conditions spatial memory performance in healthy aging

**DOI:** 10.1101/2022.10.05.510774

**Authors:** M. Durteste, L. Van Poucke, S. Combariza, B. Benziane, J-A. Sahel, S. Ramanoël, A. Arleo

## Abstract

Memory for objects and their location is a cornerstone of adequate cognitive functioning across the lifespan. Considering that human visual perception is dependent upon the position of stimuli within the visual field, we posit that the position of objects in the environment may be a determinant aspect of mnemonic performance. In this study, a population of young and older adults completed a source monitoring task with objects presented either in the upper or lower visual field. Using multinomial processing tree modeling, we revealed that while item memory remained intact in older age, spatial memory was impaired but only for objects that had been presented in the upper visual field. Spatial memory is therefore conditioned by the vertical position of information. These findings put into question the traditional view that age-related spatial mnemonic deficits are primarily attributable to high-order associative dysfunctions and suggest that they could also originate from altered encoding of object attributes.

## Introduction

To guide their behavior in everyday life, humans need to understand, attend to, and interact with the visual information in the surrounding environment. In that respect, memory for objects (i.e., item memory) and their location (i.e., spatial memory) is key to preserve cognitive performance across the lifespan. Indeed, impaired item and spatial mnemonic capacities have deleterious consequences on autonomy and quality of life in healthy aging. Previous work has often reported these deficits to have their roots in alterations of high-order associative functions such as contextual binding and executive function^1–3^. We hereby propose the view that mnemonic deficits in older age could also emerge from the deficient encoding of specific object features. Indeed, item and spatial memory are complex skills that require fine-grained visual processing of objects’ various colors, textures, sizes, and positions in space. Modified sensory function in older age may thus also contribute to the decline in memory for objects and their spatial context^4,5^.

We set forth the idea that location in space may be a particularly relevant object attribute considering that human visual perception is strongly dependent upon the position of stimuli in the visual field^6^. Not only does visual performance decrease sharply when moving away from the center of vision towards the periphery, it can also be modulated by isoeccentric locations around the visual field. One example of this effect is the horizontal-vertical anisotropy (HVA): at fixed eccentricities, visual performance is better along the horizontal than the vertical meridian. A second notable example is the vertical-meridian asymmetry (VMA), which reveals that visual ability differs between the upper and lower visual fields^7,8^. These asymmetries exist across a vast array of stimuli orientations, sizes, and luminance levels^9^. Visual acuity and contrast sensitivity^9,10^, temporal resolution^11^, spatial attention^12^, hue discrimination^13^ as well as motion processing^14–17^ are better performed in the lower visual field. A bias for the upper visual field is evident in experimental paradigms in which visual search^18,19^, change detection^20^, target identification^21–23^, or categorical judgements^24^ are implicated. While it is clear that the inhomogeneities around the visual field are pervasive across a variety of perceptual tasks, whether they extend to cognitive function, and more specifically to item and spatial memory, remains poorly understood. A couple of studies showed that the HVA and VMA could in fact influence visual short-term memory^25,26^. However, these studies used non-naturalistic stimuli in the form of gratings or blocks, and they only tested very short-term memory. More evidence is needed to test the possibility that spatial location could condition the precision with which the memory trace is encoded or retrieved.

Importantly, these vertical performance asymmetries evolve across the lifespan. A burgeoning field of research is looking into how upper-lower visual field asymmetries emerge during childhood. In a recent study, Carrasco and colleagues^27^ (2022) compared the VMA for contrast sensitivity between children and young adults. They showed that the VMA is a malleable property of the visual system that settles in beyond childhood. Another recent study concluded that perceptual asymmetries in infants reflect adaptations to typical spatial locations in the surrounding environment^28^. With regards to aging, data pertaining to upper-lower asymmetries are scarce and they remain largely inconclusive. While some research has reported that the VMA persists in late adulthood^29,30^, other psychophysical and visual search studies have revealed that vertical performance asymmetries are in fact modified in older age^31–33^.

Here, we sought to determine whether changes in visual encoding in one vertical hemifield could modulate the mnemonic performance of young and older adults. The aim of the present study was threefold; first, it strived to characterize the influence of the vertical position of information on item and spatial memory; second, it attempted to compare such visual preferences between young and healthy older adult populations; and third, it verified whether the visual field area of participants’ upper and lower visual fields could explain performance differences. To address these questions, we used a source monitoring task that assessed item and spatial memory separately and we performed kinetic perimetry to quantify the extent of individual visual fields. We computed the probabilities of remembering objects and their spatial location as a function of upper or lower visual field presentation positions using multinomial processing tree (MPT) modeling. This approach has the advantage that it renders explicit the assumptions about the relationship between item and spatial memory. Based on the few existing studies, we hypothesized that the vertical position of information would play a role in older adults’ mnemonic abilities.

## Results

A population of 52 subjects (26 young and 26 older adults) took part in this study with 6 older adults being subsequently excluded (see Methods section for more details). Participants were screened for cognitive impairment, to ensure that they were within the age-related normative ranges for short-term, long-term, and spatial memory. They all completed a battery of visual tests, including visual acuity, contrast sensitivity and kinetic perimetry before performing a desktop-based source monitoring task (Fig. 1). The experiment was divided into eight blocks that each comprised an encoding phase and a test phase. During the encoding phase, subjects had to maintain fixation on a central cross while thirty objects were presented successively in the upper or lower part of the screen. During the test phase, there were twenty objects that had been presented during the encoding phase and ten new objects shown one at a time in the center of the screen. Upon item presentation, participants had to decide if they had seen the object previously or not (i.e., *item memory*). If subjects answered that they had seen the object, they then had to decide whether the object had been presented in their upper or lower visual field (i.e., *spatial memory*).

**Figure 1.**
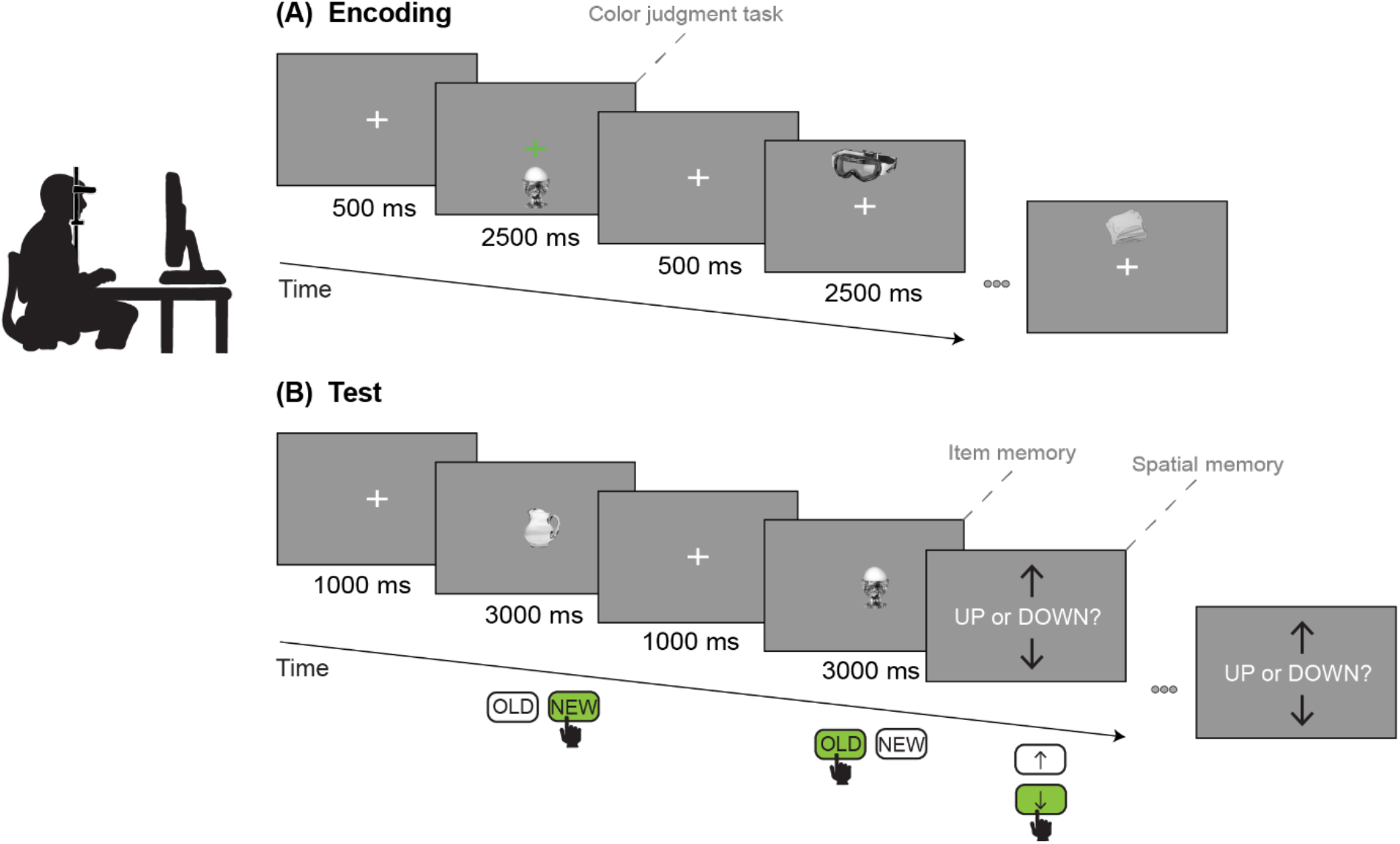
Schematic representation of the experimental paradigm. Participants performed a desktop-based source monitoring task while their gaze was recorded with an eye tracker (see Methods section for more details).

We analyzed participants’ responses using MPT modeling^34^, which can be understood in terms of binary branching trees (Fig. 2A). Each branch represents a specific cognitive process and is associated with a parameter that measures the probability of its occurrence. Terminal nodes at the end of the branches represent the observed response categories. The parameters I_up_ and I_down_ relate to item memory, S_up_ and S_down_ relate to spatial memory, and the parameters o and g measure guessing rates (see Methods section for more details).

**Figure 2.**
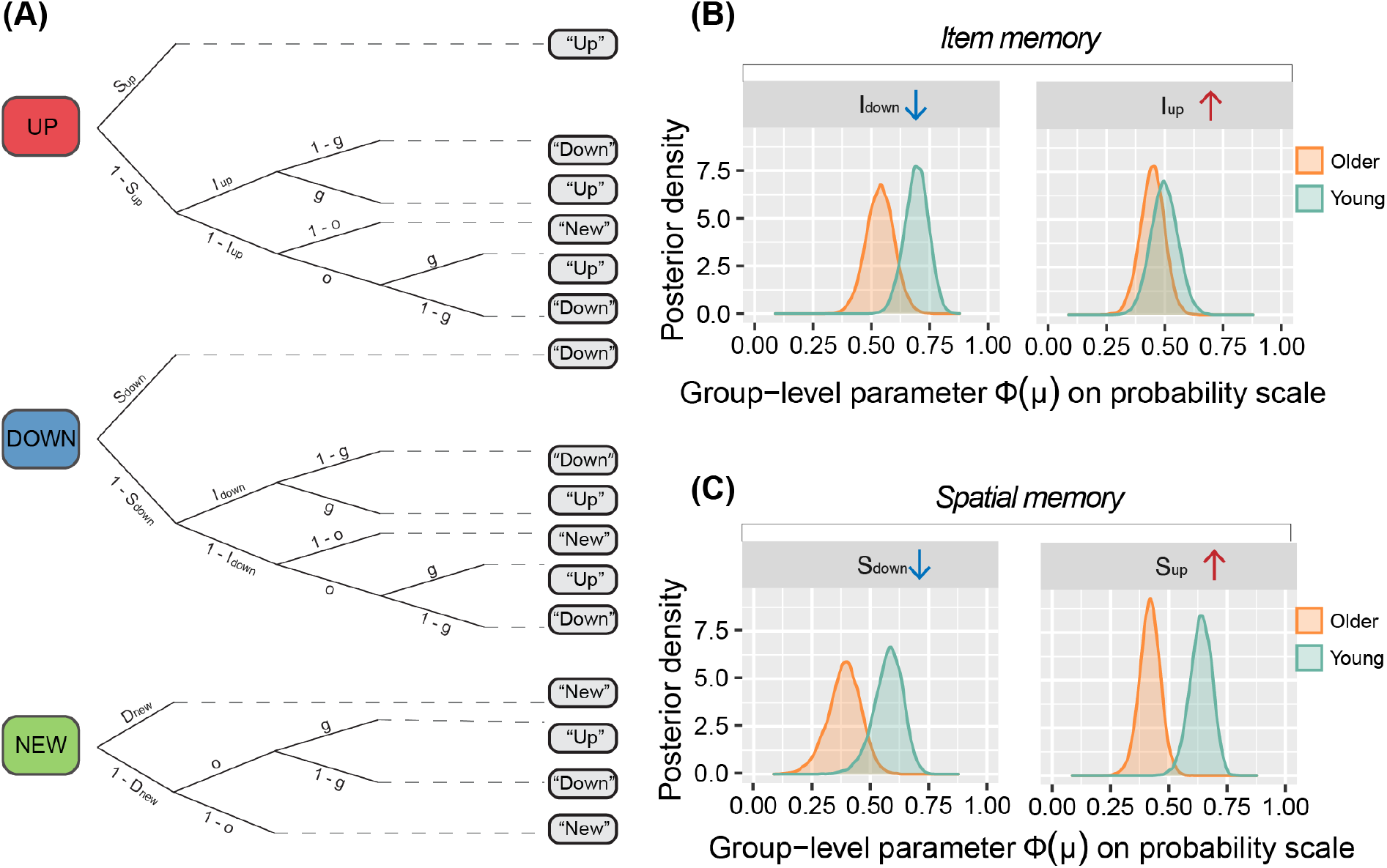
Results from the MPT analysis. **(A)** Graphical representation of the MPT model used to analyze the experimental data. **(B)** Group-level posterior distributions for the parameters related to item memory I_down_, and I_up_. **(C)** Group-level posterior distributions for the parameters related to spatial memory S_down_, and S_up_. Colored rounded rectangles represent the three different trial types: the object was presented in the upper part of the screen (“UP”), in the lower part of the screen (“DOWN”) or was not presented (“NEW”). Grey rounded rectangles represent participants’ possible answers.

We first evaluated model fits with posterior predictive *p*-values. The MPT model provided adequate fit to the data in terms of mean frequencies and covariance in both young adults (*p*_T1_ = 0.51, *p*_T2_ = 0.44) and older adults (*p*_T1_ = 0.53, *p*_T2_ = 0.58). We noted that the two parameters related to item memory (I_down_ and I_up_), the two parameters related to spatial memory (S_up_ and S_down_), and the two guessing parameters (g and *o*) were above 0 in young and older adults (Figs. 2B,C & Fig. S1). To examine differences in scores between young and older participants, we subtracted the posterior distributions of each parameter of the older group from the young group (Fig. S2).

### Item memory is invariant to the vertical position of objects

We studied the two parameters related to item memory: I_down_ and I_up_ (i.e., the probability of remembering an object that was presented in the lower and upper part of the screen, respectively). We observed that neither I_down_ nor I_up_ significantly differed between age groups (ΔI_down_ = 0.156, 95% Bayesian Confidence Interval, BCI [-0.001, 0.306]; ΔI_up_ = 0.052, 95% BCI [-0.099, 0.209]; Fig. 2B). Indeed, the parameters did not differ between young and older participants as their 95% BCI included zero. Therefore, the probability of having a sense of familiarity for an object, regardless of its position in space, was equivalent between young and older adults. We found that the probability of correctly guessing that an item was presented during the encoding phase was significantly lower in older adults (Δo = -0.064, 95% BCI [- 0.120, -0.020]; Fig. S1). In addition, despite the absence of age-related differences in object recognition, we found a significant main effect of age group (χ^2^(1) = 24.31, *p* < 0.0001) on reaction times during the item memory task (Fig. 3A). This result suggests that older adults were slower (M = 1.28s, SD = 0.46s) to respond than young adults (M = 1.01s, SD = 0.44s). We also observed a main effect of object position (χ^2^(1) = 5.69, *p* = 0.017): all participants were slightly slower to react to objects that had appeared in their upper visual field (M = 1.14s, SD = 0.47s) than in their lower visual field (M = 1.12s, SD = 0.47s). Finally, we showed that the block number had an impact on reaction times (χ^2^(7) = 343.77, *p* < 0.0001), suggesting that both young and older participants improved their performance throughout the experiment (Fig. S3).

**Figure 3.**
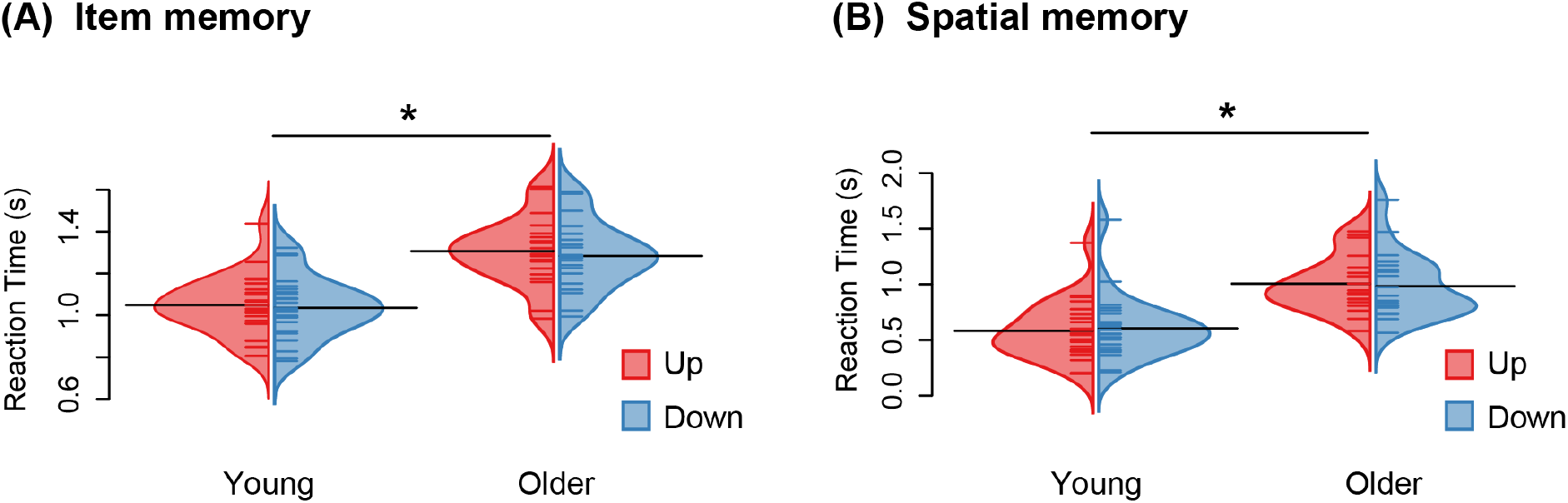
Reaction times during the **(A)** item and **(B)** spatial memory tasks as assessed by gamma generalized linear mixed models. Horizontal colored lines correspond to individual data points. Horizontal black lines correspond to the mean in each group. \**p* < 0.05

### The vertical position of objects conditions spatial memory in healthy aging

We next studied the variables related to spatial memory: S_down_ and S_up_ (i.e., the probability of remembering the position of an object that was presented in the lower and upper part of the screen, respectively). We found that while S_down_ was equivalent between age groups (ΔS_down_ = 0.188, 95% BCI [-0.002, 0.378]), S_up_ differed significantly between young and older adults (ΔS_up_ = 0.218, 95% BCI [0.086, 0.342]; Fig. 2C). These results suggest that older adults were less likely to recall the position of objects that were presented in the upper visual field. Worthy of note, the guessing parameter g, defined as the probability of correctly guessing that an item was presented in the upper visual field, was equivalent in young and older subjects (Δg = 0.080, 95% BCI [-0.079, 0.239]; Fig. S1). Moreover, we found a significant main effect of age group (χ^2^(1) = 21.83, *p* < 0.0001) on reaction times during the spatial memory task (Fig. 3B), meaning that older adults were slower (M = 1.26s, SD = 1.16s) to respond than young adults (M =, 0.69s, SD = 0.81s). Despite the performance difference reported above, there was no effect of object position on reaction times during the spatial memory task (χ^2^(1) = 0.13, *p* = 0.72). This finding suggests that participants responded equally rapidly to objects that had been presented in the upper or in the lower visual field. We also showed a significant main effect of block number (χ^2^(7) = 867.37, *p* < 0.0001) as well as a significant effect of the interaction between age group and block number on reaction times (χ^2^(7) = 39.18, *p* < 0.0001). Although all participants improved their reaction times on the spatial memory task across blocks, older adults showed a steeper learning curve (*first block*: M = 2.06s, SD = 1.58s; *last block*: M = 0.95s, SD = 0.88s) than young adults (*first block*: M = 0.99s, SD = 0.93s; *last block*: M = 0.60s, SD = 0.80s; Fig. S3).

### Measures of visual function don’t explain the age-related upper visual field deficit

We sought to verify whether the influence of the vertical position of information on spatial memory in older adults could emerge from variations in visual acuity or contrast sensitivity. There have been reports that these measures of visual function are not equivalent across the upper and lower visual fields^9,35^. Taking the latter into consideration, we conducted linear regression analyses to explore the effects of visual acuity and contrast sensitivity on the probability of remembering the position of an object that was presented in the upper part of the screen in older adults (d_up_; Fig. S4). No significant influence of visual acuity (R^2^ = 0.011, F(2, 17) = 0.099, *p* = 0.91) nor contrast sensitivity (R^2^ = 0.016, F(2, 17) = 0.14, *p* = 0.87) was found on the age-related upper visual field deficit. Next, we wondered whether the observed upper visual field deficit in older adults could arise from differences in the ratio of upper to lower visual field area as assessed by kinetic perimetry. We conducted linear mixed models to study the effects of age group, S_up_ and isopter on VMA. The interaction between S_up_ and age group did not improve the model significantly (with interaction: AIC = 2044.4; without interaction: AIC = 2043.5) and it was therefore not retained in subsequent analyses. As expected, we first found that age group had a significant impact on VMA (F(1, 40) = 6.75, *p* = 0.013; Fig. 4), with older adults having significantly higher VMA (M = 33.47, SD = 16.04) than young adults (M = 19.63, SD = 13.55). This result suggests that the upper visual field area is more impacted by the aging process than the lower visual field area. There was also a main effect of isopter on VMA (F(2, 213) = 5.90, *p* = 0.0032) with the smallest VMA corresponding to the most peripheral isopter. Finally, we didn’t find any significant effect of S_up_ on VMA (F(1, 40) = 2.26, *p* = 0.14). This result highlights that the more pronounced VMA in healthy aging is not driving the performance asymmetries in spatial memory reported in older participants.

**Figure 4.**
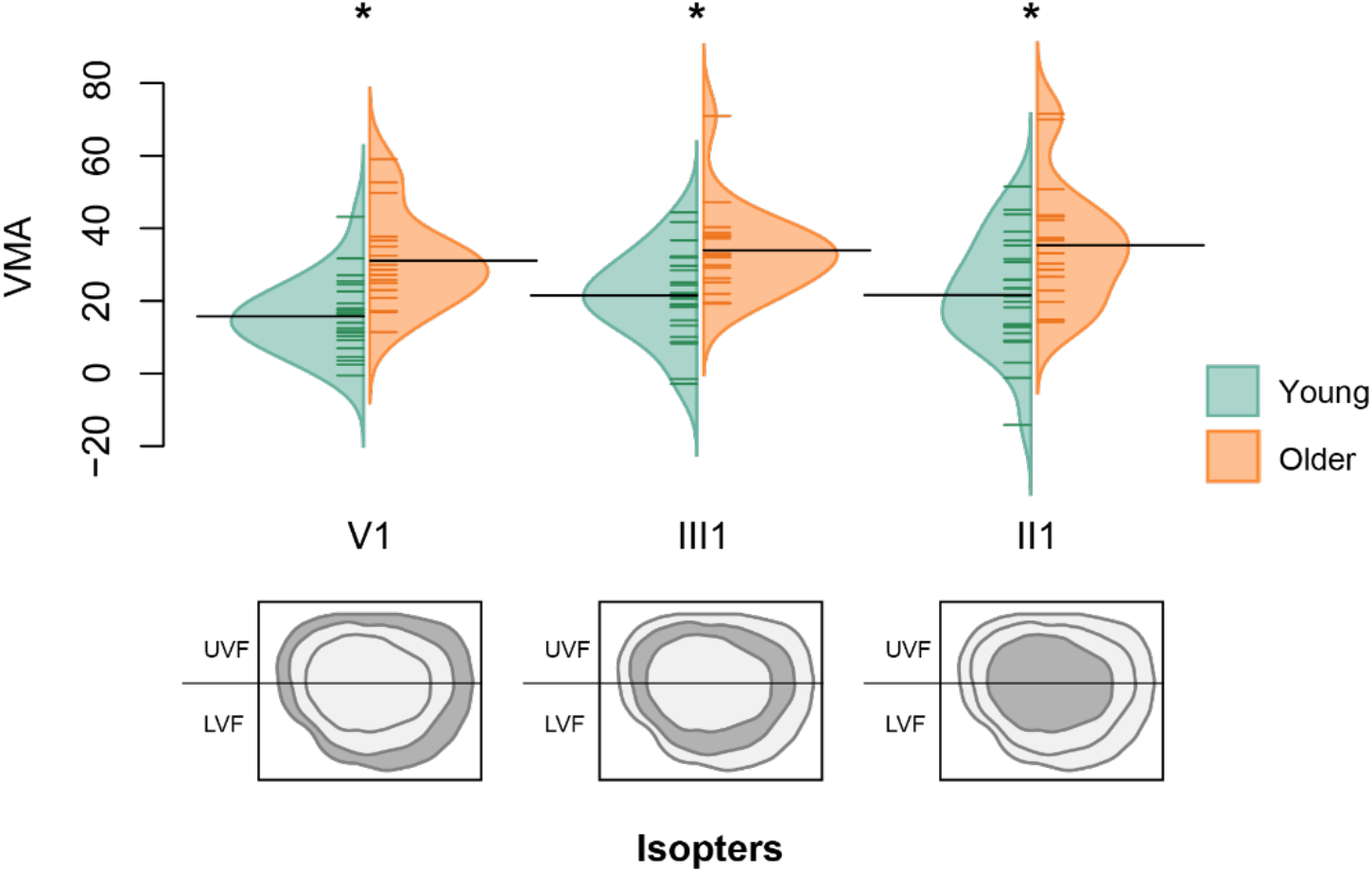
Vertical meridian asymmetry (VMA) for the visual field areas of participants across three isopters (V1, III1, II1). Visual field areas were measured with kinetic perimetry for each subject. UVF: upper visual field, LVF: lower visual field. Horizontal colored lines correspond to individual data points. Horizontal black lines correspond to the mean VMA in each group. \**p* < 0.05

Of note, we also conducted standard linear *Pr* analyses in order to compare the results with those obtained from MPT modeling. Both types of analyses led to similar conclusions. The methods and results for the standard *Pr* analyses are included in the Extended Data.

### Discussion

The aim of this study was to test the hypothesis that the visual encoding of object attributes plays an essential role in the maintenance of spatial cognition across the lifespan. More specifically, we examined whether the vertical position of visual information can impact item and spatial memory and if it does so differentially in young and healthy older adults. The findings revealed an age-related decline in spatial memory only for objects presented in the upper visual field. This result was not found to be explained by the visual field area asymmetries of participants, suggesting that this specific spatial memory deficit may stem from higher visual regions.

In accordance with the existing literature, we observed that spatial memory was impaired in older adults but that item memory showed no such age-related difference^36–39^. It is well documented that healthy aging is accompanied by extensive changes in object-location binding that are usually more pronounced than changes in simple item recognition^40^. The prominent *associative memory* theory claims that age-related deficits in episodic memory emanate primarily from inadequate associations between items and contextual information^38,41–43^. A long line of research asserts that familiarity and gist processing are the aspects of item memory that remain intact in older age^38,44,45^. The *Source-Item* model of memory that was used to explain the present data defines item memory as the probability of remembering an item given that spatial position has not been recalled. One can argue that such a process is more akin to familiarity (i.e., a global measure of memory strength) than to a detailed recollection of the object. Our study therefore confirms that the general memory of an object is preserved in healthy aging. However, it provides nuance to the widespread view that older adults are impaired at recalling spatial contextual information. We indeed revealed that older adults have significantly worse spatial memory than young adults but only for objects that were presented in the upper visual field. In other words, spatial memory may be preserved in aging for visual information situated in the lower field. Two previous studies reported an age-related loss of function in the upper visual field for rapid stimuli detection^31,32^. We extend their work to high-level cognitive functions, and we show for the first time that in older age the mnemonic trace of an object is dependent upon its vertical position in space.

Importantly, the position of information influenced spatial memory but not item recognition in healthy aging. Such a result strongly implies that the observed upper visual field deficit in spatial memory doesn’t arise from a biased distribution of covert attentional processes in older adults. Agreeing with this conclusion, accumulating evidence postulates that performance asymmetries across the vertical visual fields are not a product of attentional differences^8,46,47^. Moreover, the fact that item recognition is not influenced by the location of objects speaks to the spatial nature of vertical performance asymmetries in aging. The asymmetries appeared in older subjects only when the spatial properties of objects were solicited. Along the same lines, Genzano and colleagues (2001) concluded that upper-lower performance asymmetries were related to the visuospatial features of a stimulus^25^.

The underpinning biological basis for the reported upper visual field spatial memory deficit in older adults is unknown but it could be located anywhere along the visual pathway, from the retina to higher neural structures. A recent computational model highlighted that the VMA for contrast sensitivity was only weakly attributable to retinal factors including optics and cone sampling^48^. Furthermore, we excluded the possibility that low-level perceptual differences hindered the ability of older adults to create a detailed mnemonic trace of the object and its position. Indeed, we showed that neither visual acuity, contrast sensitivity, nor the VMA for visual field areas were associated with the probability of remembering the position of an object that was presented in the upper visual field in older participants. The VMA for visual field areas reflects the sensitivity threshold to dots of various sizes and light intensities but it doesn’t provide information relative to how that signal is processed. Notably, cortical inhomogeneities in V1 have been shown to covary with spatial frequency and contrast sensitivity behavioral asymmetries across the vertical visual field^35,49^. It is therefore likely that top-down influences from higher visual regions play a role in the age-related spatial memory decline that is specific to the upper visual field. Visual information is propagated from early visual areas through the ventral and dorsal streams that bear distinct proportions of upper and lower visual field afferents^50^. The ventral stream benefits from a larger upper visual field representation and it may be involved in far space object identification, whereas the dorsal stream relies more strongly on lower visual field inputs and it could play a role in near space visuomotor processing^51^. Importantly, regions upward of the ventral stream are closely associated with pattern separation, an essential ability to reduce the overlap between similar object locations^52^. Therefore, one possibility for the upper visual field deficit in spatial memory is an age-related neural loss that is specific to regions upward of the ventral stream. This study lends credence to the idea that the visual field biases contained in higher visual regions could have specific behavioral correlates in older adult populations.

In conclusion, our main finding that the vertical position of objects determines spatial memory performance in healthy aging strongly conveys the importance of object feature encoding in the context of cognitive decline across the lifespan. The risk of falling, the stooped posture, and the careful avoidance of obstacles on the ground are only a few examples of physical and perceptual modifications in aging that could reshape older adults’ use of visual space. Our findings also have far-reaching implications for complex cognitive skills that require adequate memory of the location of objects, such as spatial navigation or driving. In such contexts, we argue that older adults could benefit from using cues that are situated in the lower portion of their visual field or from actively training their upper visual field. Moreover, to maintain older adults’ autonomy and mobility, age-friendly environmental adaptations are to be made in line with the vertical visual field preferences hereby documented. In the future, research efforts should be committed to uncovering the neurobiological factors that link age-related visual, behavioral, and cognitive impairments.

## Methods

### Participants

We recruited a sample of 52 subjects (twenty-six young and twenty-six older adults) from the SilverSight cohort^53^ to participate in the experiment. The cohort study consists of healthy subjects without any history of psychiatric or neurological disorders who have undergone an exhaustive battery of clinical (ophthalmological, auditory, vestibular) and neuropsychological tests. All participants showed normal performance on the Mini-Mental State Examination^54^, the perspective-taking test^55^, the Corsi block-tapping task^56^, and the 3D mental rotation test^57^ (Table S1). Moreover, they had normal or corrected-to-normal eyesight. Of the total 52 participants, we excluded 6 older adults due to poor eye tracking performance (see the “Eye tracking” section for more details). Consequently, the analyses included a group of 26 young subjects (mean age: 29.2 ± 4.2 years old; 15 females and 11 males) and a group of 20 older subjects (mean age: 75.5 ± 3.7 years old; 12 females and 8 males). The Ethical Committee “CPP Ile de France V” (ID_RCB 2015-A01094-45, CPP N: 16122) approved the experimental procedures and all participants provided their written informed consent.

### Apparatus and setting

We used PsychoPy v2020.2.10^58^ on a Dell monitor with a 1280 × 1024-pixel resolution. The screen subtended 29° of visual angle in height and 36° of visual angle in width. Participants sat 57 cm from the monitor with their head positioned on the chinrest of a head-mounted monocular eye tracker (EyeLink 1000 Tower Mount, SR Research Ltd., Canada). We collected responses with a mini numeric keypad (KKmoon) positioned flat on a table in front of subjects. Young and older participants with visual corrections removed their glasses to perform the task as multifocal glasses could have biased performance in the upper or lower visual fields. Finally, the experiment took place in a quiet dim-lit room at subjects’ preferred time of day.

### Stimuli

Stimuli consisted of 327 photographs of objects selected from the datasets of the mnemonic similarity task^59^ (available at https://github.com/celstark/MST) and from the “Massive Memory” unique object image set^60^ (available at http://konklab.fas.harvard.edu/#). For this experiment, we removed images of animals or people, of objects that were strongly associated with the ground, the ceiling, or the sky (e.g., plane, chandelier) and of uncommon objects (e.g., chisel, perimeter). We resized the selected images to a 400 × 400-pixel resolution and converted them to greyscale. Then, we normalized the images to obtain a mean luminance equal to 128 on a grey-level scale ranging from 0 to 255. The objects subtended approximately 4° of visual angle in height. Figure 1 presents example images of objects.

### Procedure and task design

The experimental session began with a 9-point eye tracking calibration. The eye tracker sampled the position of the dominant eye at 1000 Hz throughout the entire session. We repeated the calibration halfway through the experiment. A 1.5-minute practice run and 8 experimental runs each lasting approximately 4 minutes structured the experimental session. Participants repeated the practice run until they understood the instructions and felt comfortable with the keypad. After the completion of 4 runs, participants had a 5-minute break during which they could remove their head from the chinrest. The session ended with a questionnaire about participants’ strategy and subjective judgement of task difficulty.

We adapted the design of the paradigm from the classic source monitoring paradigm that assesses both item recognition and source memory^61^. Each run comprised an encoding phase and a test phase (Fig. 1). The encoding phase started only once participants had stable fixation on the central cross for more than 800 ms. The task presented 30 unique objects sequentially for 2500 ms in the upper or lower part of the screen along the vertical meridian. A 500 ms interval separated each stimulus presentation. Participants had to maintain fixation on the central cross throughout the entire duration of the encoding phase. To facilitate fixation, participants performed an additional task that consisted of pressing a button when the cross turned green. During the test phase, participants saw 20 objects that had been presented during the encoding phase and 10 new objects (i.e., distractors), ordered pseudo-randomly. Stimuli stayed in the center of the screen for 3000 ms and subjects decided whether the object was old or new by pressing the appropriate key (item memory). If participants responded “Old”, we subsequently tested their memory of the position of the object (spatial memory). Participants determined whether the object had appeared “up” or “down” in a self-paced manner. They didn’t receive any feedback. A 1000 ms interval separated each trial of the test phase to leave enough time for older adults to reposition their fingers on the keypad. All timings were decided based on previous studies using the source monitoring paradigm and on pilot testing with older participants.

To minimize order effects, we counterbalanced the sequence of runs using a Latin square design. We also counterbalanced the studied objects by size and category across screen positions and across runs. Finally, we counterbalanced the position of objects across participants. One half of participants saw *object 1* in the upper part of the screen and the other half saw *object 1* in the lower part of the screen.

### Visual testing

Trained orthoptists performed all visual testing. We assessed visual acuity and contrast sensitivity on all participants using the LogMAR scale and the Pelli-Robson score, respectively. We also performed kinetic perimetry on the Octopus 900 (Haag-Streit, Switzerland). We obtained participants’ visual fields from both eyes using stimulus sizes of V1e, III1e, II1e, and I1e at 4 degrees per second for isopter charting. We adjusted participants’ responses to stimuli presentation for reaction times. We then used custom MATLAB code to process the visual field data for the left and right eyes of each participant. The latter extracted upper and lower visual areas along the horizontal meridian for the isopters corresponding to stimulus sizes of V1e, III1e, and II1e. We subsequently computed the vertical meridian asymmetry index as the difference in area between the lower and upper visual fields, divided by the mean of the two areas, and multiplied by 100:

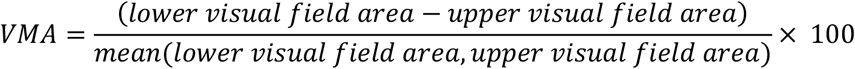

Indices that are close to 0 indicate no asymmetry between the areas of the two hemifields. Higher indices, on the other hand, reveal asymmetry with a larger area in the lower visual field than in the upper visual field. Note that we excluded two older subjects from the visual field analyses as their kinetic perimetry data were not exploitable.

### Data analysis

We performed all data analyses using R version 4.0.3 in RStudio version 1.4.1103 (R Core Team, 2020; RStudio Team, 2021). We used the interquartile range criterion to identify outliers in the data. Two young participants qualified as outliers statistically. However, removing them from the analyses described below did not change the results meaningfully and we kept them in.

#### Eye tracking

To classify gaze data into fixations, we chose the velocity-based algorithm implemented in the *saccades* R library^62^. Prior to analysis, we removed all fixations that lasted under 100 ms. We then used the 2/4 method and corresponding Fujii classification to assess fixation stability and identify participants who failed to maintain central fixation throughout the encoding phases^63,64^. The latter method considers participants to have unstable fixation if less than 75% of their fixation points are outside a 4° circle. As previously mentioned, we consequently excluded six older adults.

#### Reaction times

Reaction time is positively skewed by nature. Following recent recommendations on how best to model such data^65^, we conducted two gamma generalized linear mixed models with a log link function to analyze reaction times for the item memory and spatial memory tasks. In both models, the random-effects structure included subject and block as random intercepts along with the random effect of trial in each age group. The fixed effects were age group, object position, and block number, sex was removed as it did not improve model fit.

#### Multinomial processing tree (MPT) modeling

We performed MPT modeling to distinguish item and spatial memory between objects that were presented in the upper part of the screen and objects that were presented in the lower part of the screen. By analyzing response frequencies, MPT models provide probability estimates of latent cognitive processes implicated in a specific paradigm^66,67^. Indeed, MPT models rely on extensive theoretical work making them ideally suited to disentangle the meaningful psychological phenomenon implicated in the task at hand. In the source monitoring paradigm spatial memory indices are often biased by item recognition and guessing prevalence, MPT measurements are therefore particularly useful as they precisely isolate spatial memory^34,68^. The *Item-Source* model in which spatial memory is considered to be dependent upon item recognition has been the model of choice in most studies using MPT modeling for source monitoring tasks. However, Cooper and colleagues^69^ (2017) recently put into question the theoretical assumptions behind this unidimensional account of memory. The dual-process *Source-Item* model presents itself as an interesting alternative by suggesting that both spatial and item memory can be supported by a recollection mechanism while item memory can also be subtended by familiarity alone when recollection fails^61,70^. Taking the latter into account, we chose to apply the *Source-Item* model to the present paradigm. According to the *Source-Item* model, I_up_ and I_down_ depict the phenomenon of familiarity that can occur when recollection fails for objects presented in the upper or lower visual fields. On the other hand, S_up_ and S_down_ reflect the process of recollection in which both the object and its position are remembered at once for objects that were presented in the upper or lower visual fields. The parameter o corresponds to the probability of correctly guessing that the object was presented, and the parameter g estimates the probability of correctly guessing the position of an object that was presented in the upper part of the screen, given that item recognition has failed. We fit the behavioral data to the MPT model with the *TreeBugs* package in R^71^. This R package allows latent-trait models to be fit using Just Another Gibbs Sampler (JAGS) to estimate posterior distributions. Importantly, latent-trait hierarchical models take into account interindividual variability^72^. We kept the weakly informative default priors provided by the *TreeBugs* package and we included sex as a covariate in the MPT models. The latter estimate the latent-trait parameters using Markov chain Monte-Carlo (MCMC) simulations in JAGS. The algorithm performed 50,000 iterations with a burn-in period of 10,000 samples and thinning factor of 10. R < 1.05 indicated that a parameter had reached good convergence. We relied on posterior predictive p-values p_T1_ and p_T2_ to evaluate the goodness-of-fit of the final models^72^. The p_T1_ and p_T2_ variables measure model fits in terms of means and covariance, respectively. Predictive p-values > 0.05 indicate that the data is well accounted for by the model. We also inspected the Watanabe-Akaike information criterion (WAIC) as it quantifies relative model fits by subtracting a correction for the number of parameters included.

## Acknowledgments

The authors would like to express their most sincere gratitude to the volunteers who took part in this study. We also thank Elisa Tartaglia and Valérie Parmentier (Essilor International) for their helpful inputs.

## Author Contributions

Study design: MD, SR, AA; Data acquisition: MD, LVP, SC; Data processing: MD, BB; Manuscript writing: MD, SR, JAS, AA.

## Supplementary information

**Table S1.** Summary of young and older participants’ scores on various cognitive tasks. M: males; F: females; SD: standard deviation; MMSE: mini mental state examination.

|  | Groups |  |
| --- | --- | --- |
|  | Young adults<br>11 M / 15 F | Older adults<br>8 M / 12 F |
| | Mean (± SD) | Mean (± SD) |
| MMSE | 29.2 (± 1.1) | 28.2 (± 1.5) |
| 3D Mental Rotation | 15.9 (± 5.7) | 8.5 (± 5.3) |
| Corsi forward | 6.3 (± 1.1) | 4.5 (± 0.8) |
| Corsi backward | 6.0 (± 1.2) | 4.5 (± 0.8) |
| Perspective taking | 23.8 (± 19.1) | 51.2 (± 28.2) |

**Figure S1.**
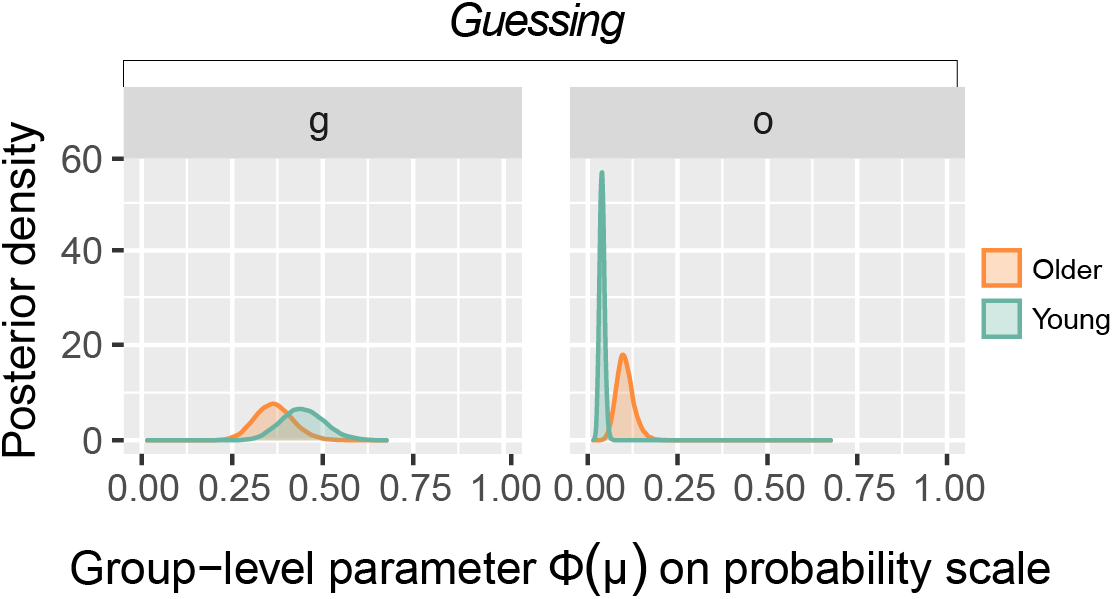
Results from the MPT analyses. Group-level posterior distributions for the parameters related to guessing rates g and o. The parameter o describes the probability of correctly guessing that the item was presented. The parameter g on the other hand describes the probability of correctly guessing that the item was presented in the upper visual field.

**Figure S2.**
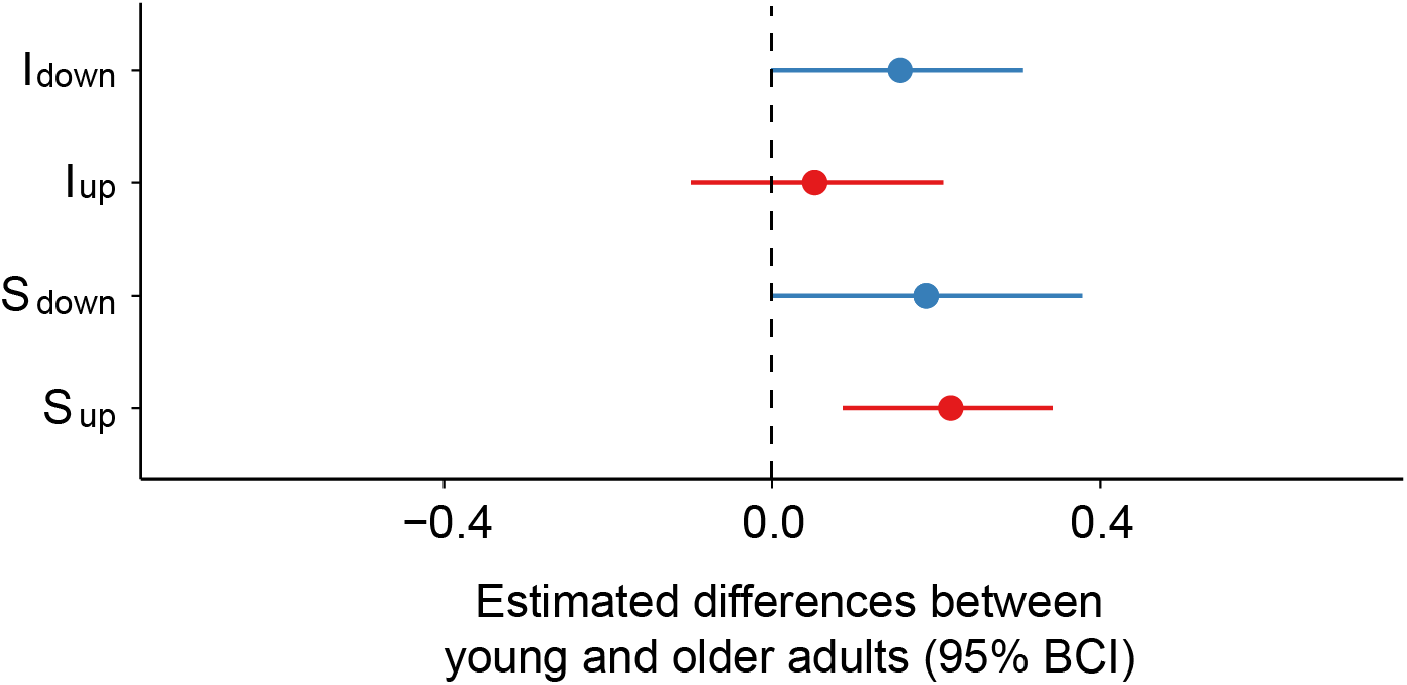
Differences in parameter estimates between young and older adults. The dot represents the parameter estimate and the line is the 95% Bayesian credible interval (95% BCI). Parameters in blue are those linked to the lower visual field while parameters in red are those linked to the upper visual field.

**Figure S3.**
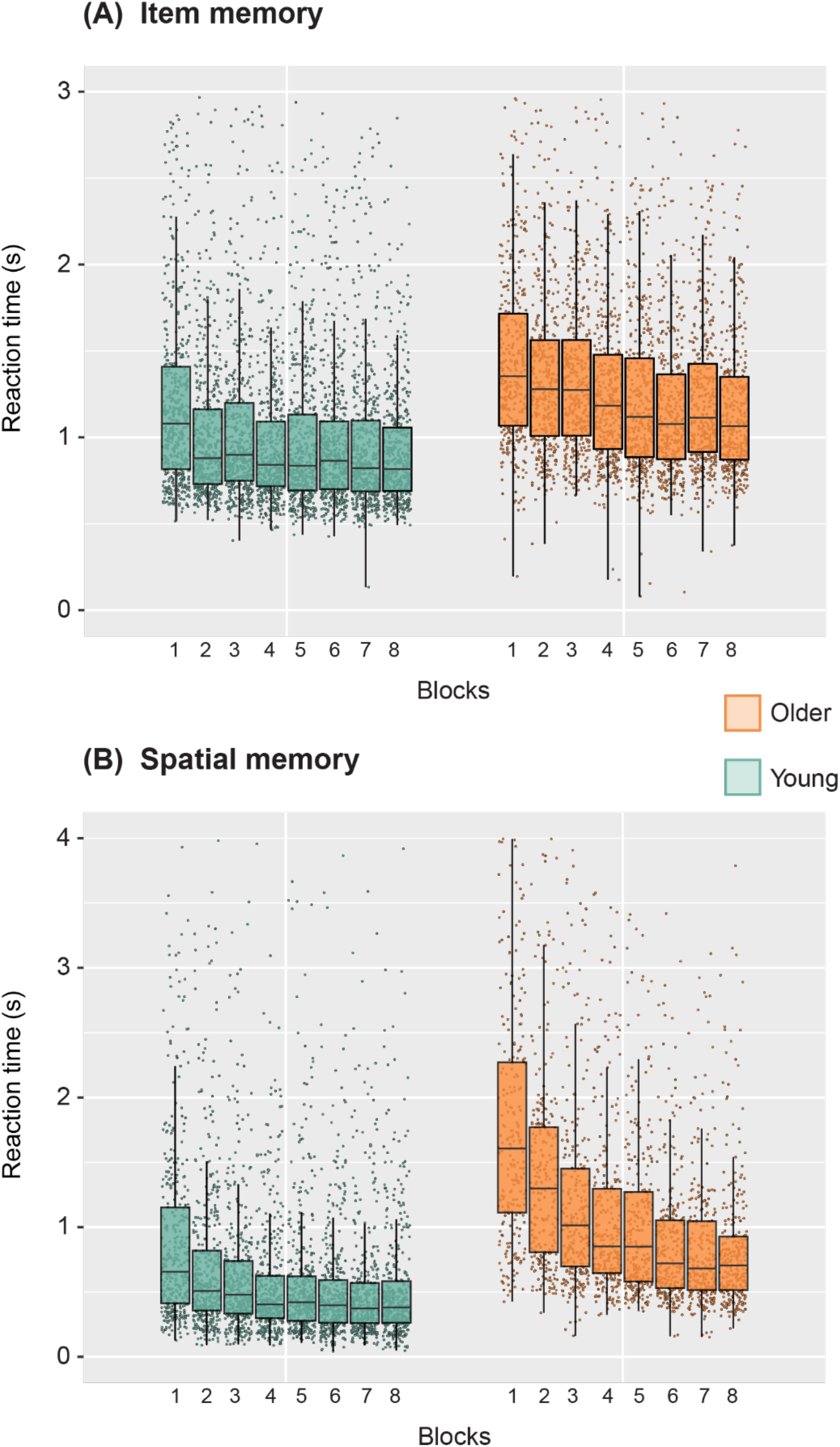
Reaction times (seconds) across blocks for the **(A)** item memory and the **(B)** spatial memory task. Each dot is a unique trial.

**Figure S4.**
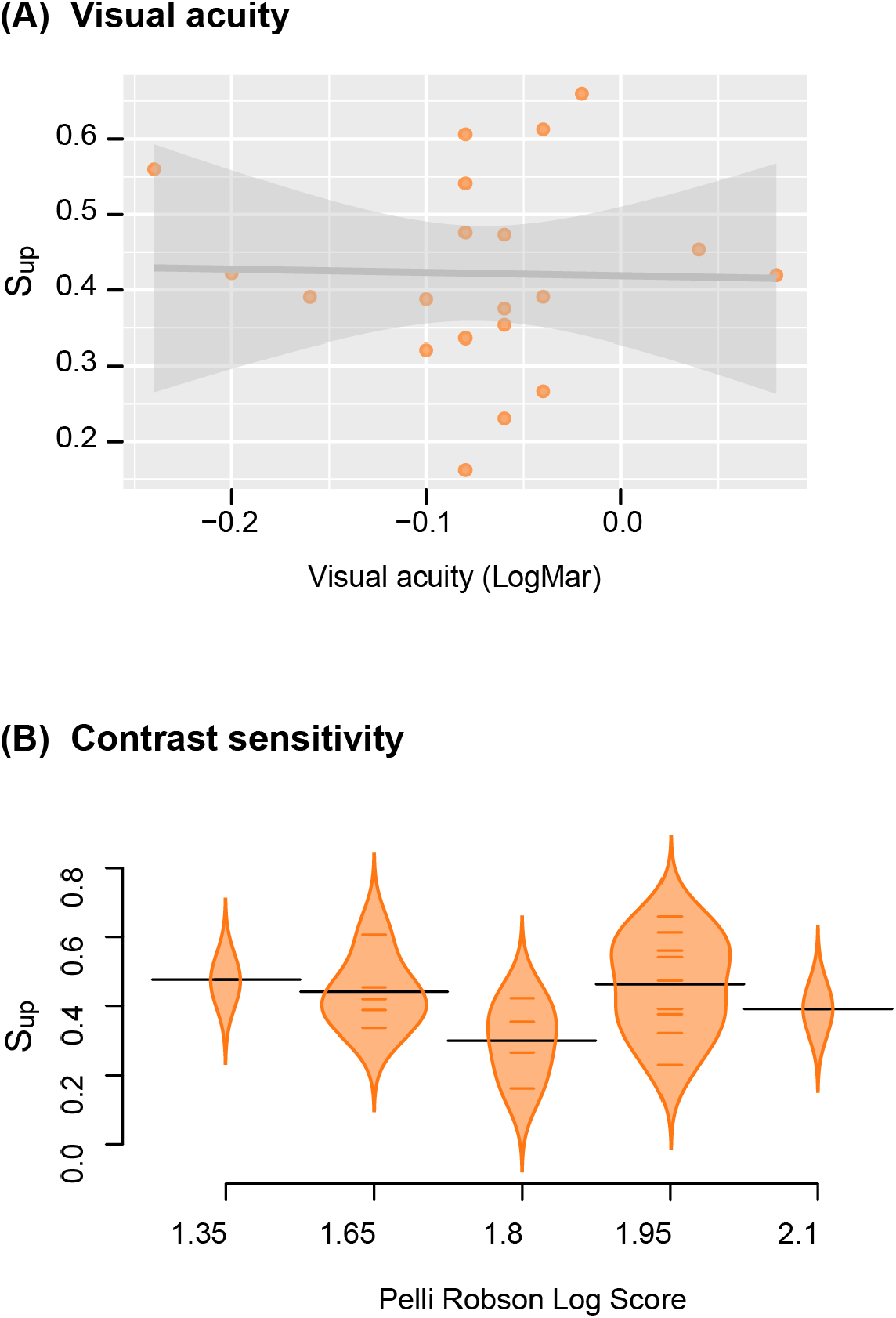
Associations between spatial memory performance and measures of visual function in older adults. **(A)** The probability of remembering the position of an object that was presented in the upper part of the screen (S_up_) as a function of visual acuity assessed with the LogMAR scale. **(B)** The probability of remembering the position of an object that was presented in the upper part of the screen (S_up_) across various Pelli-Robson contrast sensitivity scores. Horizontal colored lines correspond to individual data points. Horizontal black lines correspond to the mean.

